# Wastewater-impacted Skagerrak Sea microbiomes anaerobically demethylate micropollutants

**DOI:** 10.64898/2026.01.06.697959

**Authors:** Tetyana Gilevska, Amelia-Elena Rotaru, Konstantinos Anestis, Alexis Fonseca, Steffen Kümmel, Martin Krauss, Pedro A. Inostroza, Stefano Bonaglia

## Abstract

Methylated micropollutants such as naproxen and caffeine persist in wastewater effluents and accumulate in coastal sediments, including Hakefjorden, Skagerrak Sea, yet their anaerobic fate and role in methane emissions remain unresolved. Here we traced the fate and microbiome responses to ¹³C-labeled naproxen and caffeine in sediment microcosms. We show that naproxen underwent rapid O-demethylation to desmethylnaproxen, with 90% ± 15.5% removed within 25 days, producing primarily ¹³CO₂ and some ¹³CH₄. Naproxen enriched methylotrophic and hydrogenotrophic *Methanomicrobia*, alongside *Lokiarchaeia, Bathyarchaeia*, and bacterial taxa like *Eubacterium* and *Syntrophomonadaceae*. Metagenomics revealed O-demethylation genes in enriched bacterial MAGs affiliated with uncultured *Thermoanaerobaculia*, indicating a bacterial demethylation potential. In contrast, caffeine was largely recalcitrant to degradation (∼85% ± 5% remaining), yet its ^13^C-labeled N-methyl groups fueled ¹³CH₄ production, coinciding with enrichment of *Bathyarchaeia* and *Methanosarcina*. These results show that methylated micropollutants can activate both bacterial and archaeal demethylation pathways in coastal sediment microbiomes.

## 1. Introduction

Thousands of pharmaceuticals, pesticides, personal care products, and industrial chemicals, among others, enter aquatic ecosystems through wastewater treatment discharge, agricultural runoff, and improper disposal^1,2^. These chemicals, collectively referred to as micropollutants, are routinely detected in freshwater and marine environments at concentrations ranging from picograms to micrograms per liter, raising significant concerns^3^ due to their role in driving the aquatic biodiversity crisis^4^.

Caffeine and naproxen, which we use as a stimulatory substance and a nonsteroidal anti-inflammatory drug, respectively, are micropollutants frequently detected in aquatic environments, where they persist despite wastewater treatment. Although treatment removal efficiencies are reported as high (64-100% for caffeine and 40-100% for naproxen), both compounds remain detectable in effluents, leading to contamination of receiving waters at ng/L to µg/L and of sediments at ng/g levels^5–7^. Owing to its ubiquity, caffeine is even used as a tracer for wastewater pollution^8^. Their persistence raises ecological concerns: caffeine disrupts biochemical and cellular processes in fish^9,10^ and is a neurotoxicant for aquatic animals^11^, while naproxen and its metabolites have been associated with endocrine disruption^12^, mutagenicity, genotoxicity, carcinogenicity, and reproductive toxicity in aquatic animals^13^.

Biodegradation and photodegradation are the primary processes governing the removal of caffeine and naproxen, with reported half-lives of 0.8 – 66 and 1.5 - 100 days, respectively^14–16^. While photodegradation is restricted to surface waters, microbial degradation occurs under both aerobic^10,13^ and anaerobic conditions^17–19^. Aerobic degradation of caffeine has been demonstrated in a broad range of bacterial families (*Pseudomonadaceae, Alcaligenaceae, Bacillaceae, Brevibacteriaceae, Enterobacteriaceae, Paenibacillaceae*, and *Xanthomonadaceae*)^20^, while aerobic degradation of naproxen has been observed in comparably fewer taxa (*Bacillaceae*, *Xanthomonadaceae*, and *Planococcaceae*)^21^. In contrast, anaerobic degradation of both compounds has only been reported in mixed microbial communities inhabiting sediments or anaerobic sludge^17–19^. The specific organisms and pathways responsible for degradation under anoxic conditions, however, remain largely unknown.

Under anoxic conditions, a critical initial step in the microbial degradation of methylated organic compounds like caffeine and naproxen is demethylation - the enzymatic cleavage of a methyl group^9,13^. This reaction often determines whether further mineralization can proceed and whether degradation products are further funneled into methanogenesis. In anaerobic digesters, naproxen demethylation is linked to methane production^18,22^, and proposed to involve synergistic interactions between acetogens, acetate oxidizers, and methanogens^18^. In marine sediments, by contrast, acetogens and fermenters are thought to mediate complete O-demethylation of naproxen under sulfate-rich conditions, although it is unclear whether this process yields methane^19^. Recent findings further demonstrate that the marine methylotrophic methanogen *Methermicoccus shengliensis* can directly demethylate O-methylated aromatic compounds to methane^23^, underscoring the potential role of methanogens in demethylation of micropollutants in the marine environment.

Resolving whether bacteria or methylotrophic archaea drive demethylation in coastal sediments is essential for predicting the environmental impact and greenhouse gas outcomes from these ubiquitous methylated pollutants^2^. In sulfate-low anoxic environments, certain bacteria operating the Wood-Ljungdahl pathway may cleave methyl groups and convert them to acetate or CO_2_, a route that yields little or no CH_4_. In sulfate-rich, anoxic environments such as marine sediments, sulfate-reducing bacteria are expected to mineralize methylated micropollutants, further suppressing methane release. If, however, methylotrophic methanogenic archaea gain access to methylated micropollutants, the methyl groups can be channeled directly into methane. Clarifying the predominant demethylating guild is therefore key to assessing whether naproxen, caffeine, and related contaminants represent a previously overlooked pathway of marine CH₄ production.

To address this knowledge gap, we targeted two ubiquitous methyl-bearing micropollutants: (i) caffeine (a common stimulant with three N-methyl groups) and (ii) the analgesic naproxen (one O-methyl group), and asked which microbial guilds demethylate these micropollutants in coastal anoxic sediments^24–26^. We incubated ¹³C-labeled caffeine and naproxen in strictly controlled microcosms prepared from wastewater-impacted Hakefjorden (Swedish West Coast) sediments and traced their degradation to intermediates and ¹³C-methyl group conversion into ¹³CH₄ and ¹³CO₂. Isotope-resolved conversion profiles were related to relative taxon abundances (16S rRNA gene amplicon sequencing data) and methyl-transfer gene inventories (metagenomics). Parallel incubations with ^13^C-acetate, ^13^C-carbonate, and ^13^C-methylamine provide benchmark taxa and pathways for acetoclastic, CO_2_-reductive and methylotrophic methanogenesis. The distinct chemistries of the two substrates shaped their fates. Naproxen’s O-methyl groups were almost completely removed, whereas only a fraction of the three N-methyl groups on caffeine’s heteroaromatic purine ring were metabolized. By pairing precise ^13^C-isotope tracing with high resolution community profiling and metagenomics, we pinpoint the anaerobic taxa responsible for these demethylation routes and evaluate their potential to channel pollutant-derived methyl carbon into methane.

## Results & Discussion

### Micropollutant profile of sampled sediments

We screened the upper 10 cm of Hakefjorden sediment, collected ∼50 m north of the Strävliden wastewater treatment plant outfall, for 183 target micropollutants. Polymer additives, personal care and household products, and pharmaceuticals were the classes with the highest representation, together accounting for ∼50% of all detected micropollutants (full list in <u>10.5281/zenodo.15268726</u>).

Micropollutant class frequencies were comparable between 0-4 cm and 4-10 cm horizons, indicating limited depth-dependent attenuation within this interval. Caffeine and naproxen were present at up to 7 ng/g and 4 ng/g, respectively, within the range reported for coastal sediments worldwide^24,26^, highlighting chronic exposure of coastal microbiomes to these methyl-bearing micropollutants. In the 0–10 cm layer used for incubations, geochemical profiles indicate sulfate-rich (22 ± 1 mM) and organic-rich (2.5 ± 0.1 %, fraction of organic carbon (*f_OC_*)) conditions with minimal methanogenesis (0.4 ± 0.3 µM CH₄). These characteristics are typical of Swedish West Coast sediments^27^.

### Degradation of micropollutants in microcosms and isotopic tracing of demethylation

Methanogenic microcosms established from the 0-10 cm horizon of the Hakefjorden sediment showed rapid naproxen removal, with 90 ± 15% removed after 25 days. This depletion was accompanied by the accumulation of the degradation intermediate, desmethylnaproxen, to 83 ± 39% of the original naproxen concentration, confirming active degradation. In contrast, caffeine was largely resistant to degradation, with 85 ± 5% still detectable after 25 days (Fig. 1A). Because both substrates were added at low concentrations (<1 mM), methane buildup in amended slurries was not anticipated to significantly exceed that of unamended controls. Instead, we traced degradation by tracking the conversion of ^13^C-labeled methyl groups (Fig. 1B). Naproxen demethylation produced stronger ^13^CO_2_ enrichment (Δδ^13^C= 53 ± 16‰) than methane (Δδ^13^C= 27 ± 3‰), consistent with bacterial O-demethylation yielding CO₂ as the primary product. In contrast, caffeine incubations showed no ^13^CO_2_ enrichment, but measurable ^13^CH_4_ enrichment (Δδ^13^C= 28 ± 16‰) (Fig. 1B), indicating that a subset of caffeine-derived methyl groups was funneled into methanogenesis despite low overall degradation. The contrasting isotopic signatures reveal distinct anaerobic demethylation pathways for naproxen and caffeine in Hakefjorden sediment slurries.

**Figure 1.**
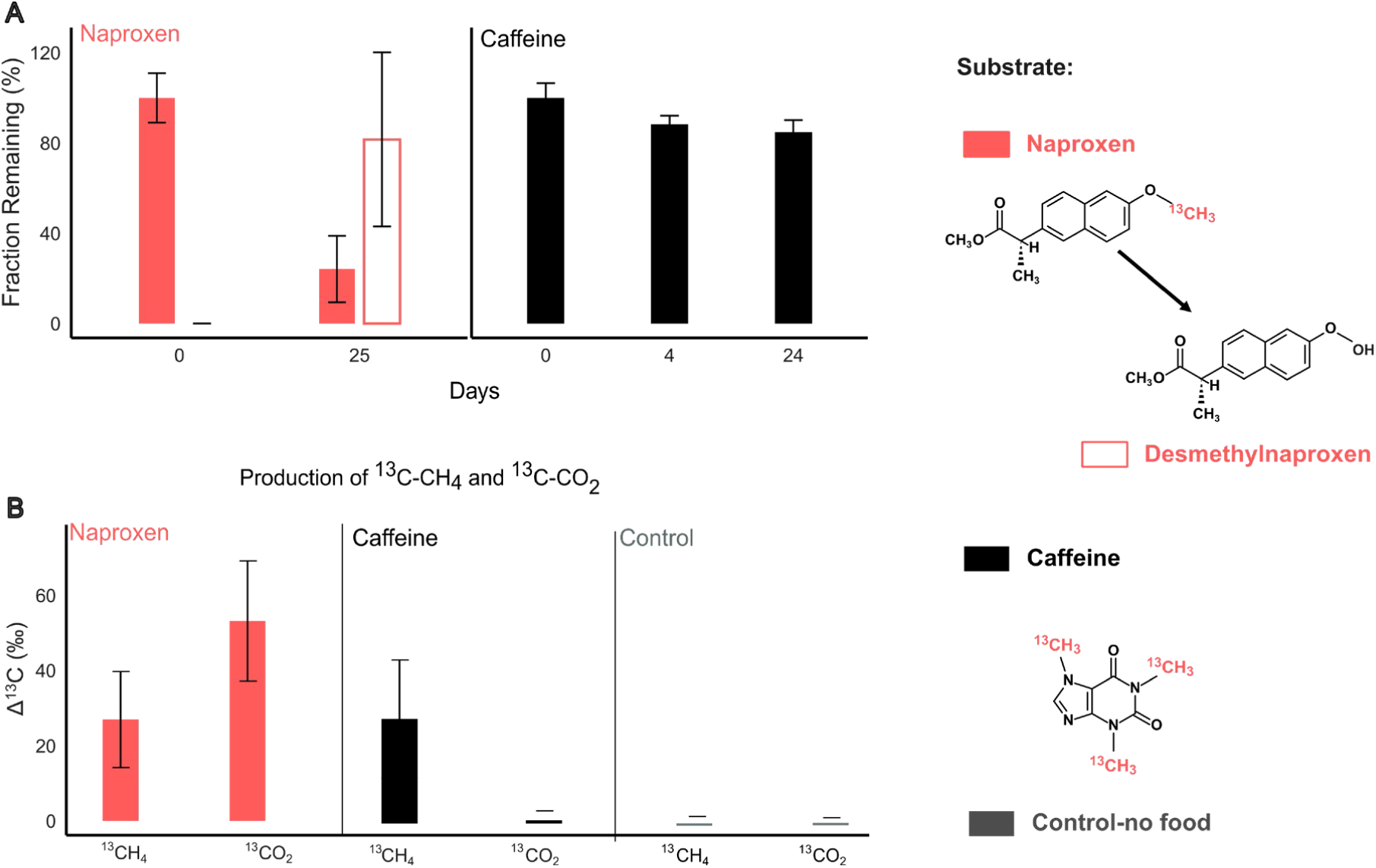
Anaerobic degradation and isotopic tracing of demethylation of two micropollutants (caffeine and naproxen) in Hakefjorden sediment slurries. (A) Rapid removal of a large fraction of naproxen after 25 days of incubation, accompanied by accumulation of its degradation product, desmethylnaproxen; (B) Isotopic enrichment of ^13^CH_4_ and ^13^CO_2_ in microcosms amended with ^13^C-naproxen and ^13^C-caffeine after 25 days. Control treatments lacked ^13^C-labeled substrates, so no enrichment of ^13^C-products could be measured.

To contextualize these results, we compared methanogenic activity on reference methanogenic substrates fueling methylotrophic, hydrogenotrophic, or acetoclastic pathways. ^13^C-methylamine was rapidly converted to ¹³CH_4_, demonstrating robust methylotrophic potential, whereas ^13^C-carbonate supported delayed hydrogenotrophic methanogenesis, noticeable only after the second headspace purge with a H_2_: CO_2_ (80:20) gas mix. ^13^C-acetate supported negligible CH_4_ production, although both ^13^C-acetate and ^13^C-methylamine incubations showed ¹³CO₂ enrichment. Sulfate concentration decreased in all microcosms, though only minimally except in the carbonate- and acetate-amended treatments. Together, these patterns confirm elevated demethylating capacity in shallow marine sediments, indicating that resident methanogens and demethylating bacteria are well-prepared to exploit methyl-bearing micropollutants.

### Archaeal and bacterial taxa with demethylation potential enriched by micropollutants Community shift on naproxen

Naproxen and caffeine exposure led to distinct shifts in microbial community composition, as revealed by 16S-rRNA gene amplicon sequencing and metagenomic analyses (Fig. 2A–D).

**Figure 2.**
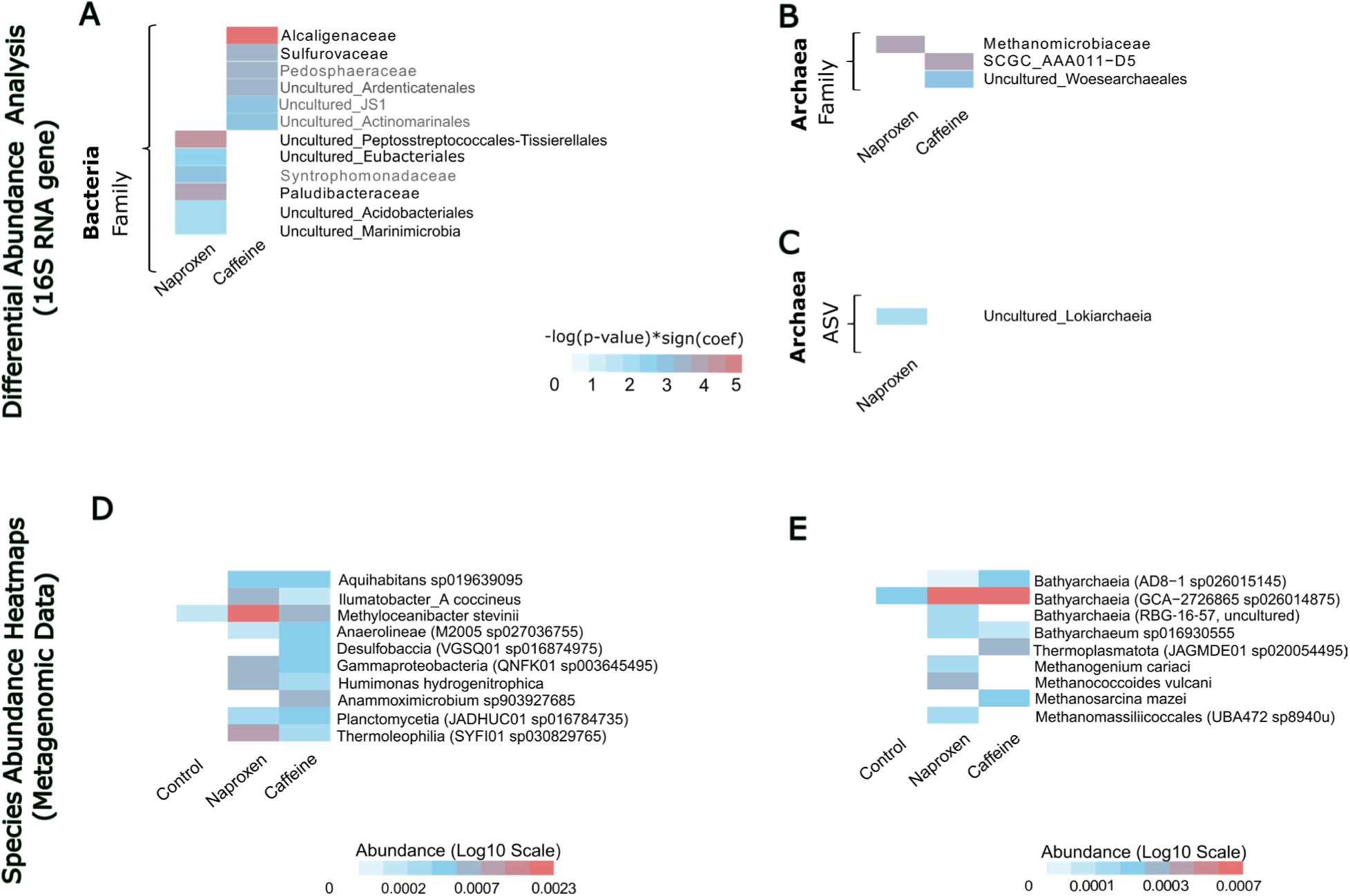
Differential relative abundance of microbiomes in caffeine and naproxen treatments relative to no-food controls. Panels (A–C) present 16S rRNA gene amplicon sequencing analyses of (A) bacterial families, (B) archaeal families, and (C) ASVs, with enrichment scores relative to unamended controls. Color intensity represents the enrichment score calculated as (–log(p-value) × sign(coef)) using MaAsLin2. Panels (D) and (E) show metagenomic species abundance heatmaps for the top five (D) bacterial and (E) archaeal species with >2-fold enrichment and relative abundances >0.01% compared to unamended controls. Taxonomic names in black font (panels A, B, and C) represent taxa detected in both amplicon and metagenomic datasets, while those in grey font indicate taxa detected only by amplicon sequencing. All taxa displayed in panels A, B, and C have adjusted p-values (FDR) smaller than 0.05 as determined by edgeR.

Naproxen selectively enriched bacterial and archaeal taxa, typically associated with O-demethylation, fermentation, syntrophy and methanogenesis.

Our ^13^C-labeling experiments demonstrated that naproxen degradation proceeds primarily via bacterial O-demethylation, yielding ^13^CO_2_ as the primary product. *Eubacteriales,* which encode the enzymatic machinery required for demethylation of methylated aromatics^28,29^ were uniquely enriched under naproxen exposure (p≤0.05, Fig. 2A). Their selective enrichment in these slurries originating from wastewater-contaminated marine sediments mirrors the response of *Eubacteriales* in communities from wastewater exposed to naproxen^18,19^ reinforcing their central role as primary demethylators of naproxen across wastewater-impacted ecosystems.

Naproxen also promoted the increase in relative abundance of multiple fermenting taxa that are likely to degrade desmethylnaproxen and other complex organics into smaller intermediates that fuel downstream syntrophs. Taxa likely involved in the desmethylnaproxen fermentation step may include *Paludibacteraceae* (p≤0.05, Fig. 2A), uncultured *Acidobacteriales* (undetected in control, enriched 33-fold p≤0.05, Fig. 2A), *Peptostreptococcales-Tissierellales* (enriched 3.4-fold, p≤0.05, Fig.2A), *Anaerolineae* (enriched 2.9-fold, Fig.2D), *Planctomycetia* (enriched 4.4-fold, Fig.2D) and the deep-sea Gammaproteobacteria *Woeseiaceae* (enriched 11.7-fold, Fig.2D). The fermentation products generated by these lineages provide substrates for syntrophic oxidation downstream.

A key syntrophic group enriched exclusively under naproxen was *Syntrophomonadaceae* (p≤0.05, Fig. 2A), classical ß-oxidizers that degrade short-chain volatile fatty acids and alcohols syntrophically^30,31^. These organisms are the primary thermodynamic bottlenecks in anaerobic organic matter degradation^32^, because their oxidation reactions are unfavorable unless their metabolic products (*e.g.,* formate & H_2_) are immediately removed. This constraint is relieved by methanogens, which efficiently scavenge these products and render syntrophic oxidation reactions thermodynamically feasible^33^.

To identify methanogens that could partner with the enriched syntrophs, we examined the archaeal taxa enriched under naproxen, but absent or low in the unamended control. The methanogen most likely to establish a H_2_-dependent partnership with *Syntrophomonadaceae* is the hydrogenotrophic *Methanomicrobiaceae* representative *Methanogenium cariaci,* whose relative abundance increased ∼2.9-fold (Fig. 2E). Consistent with this, the *Methanomicrobiaceae* family was exclusively enriched on naproxen (p≤0.05, Fig. 2B) compared to control incubations.

Because our ^13^C-labeling experiments showed that a small fraction of the ^13^C-product is ^13^CH_4_, we examined whether methanogens that can demethylate were also enriched on naproxen. Two methanogens with known O-demethylation potential were enriched: *Methanococcoides vulcanii* (enriched ∼5.9-fold) and *Methanomassiliicoccus* sp. (undetected in control)^34^. Both encode methyltransferase systems capable of O-demethylation, although neither has been implicated in O-demethylation of aromatic compounds directly^34,35^.

Beyond the methylotrophic and hydrogenotrophic methanogens, naproxen also enriched archaeal lineages assumed to be involved in fermentation and syntrophic interactions. These include uncultured *Lokiarchaeia* (undetected in control, p≤0.05, Fig. 2C) likely contributing additional fermentative capacity, and uncultured *Bathyarchaeia* (enriched ≥2.5-fold, Fig. 2E) a group known to produce H₂ and acetate during syntrophic degradation^36–38^ possibly interacting syntrophically with the hydrogenotrophic *Methanogenium*. Some *Bathyarchaeia* can even degrade lignin-derived aromatics^39^ suggesting a broader role in processing complex organic matter under naproxen exposure.

O-demethylation of aromatic compounds has so far only been demonstrated in some *Methanosarcinales* methanogens such as *Methermicoccus shengliensis*^40^. Whether methylotrophic methanogenic lineages like *Methanococcoides* or *Methanomassiliicoccus* perform similar reactions remains to be demonstrated. The predominance of ¹³CO₂ over ¹³CH₄ during naproxen demethylation, together with enrichment of syntrophic bacteria and hydrogenotrophic methanogens, suggests that bacteria such as *Eubacteriales* perform the primary O-demethylation step. At the same time, the measurable production of ¹³CH₄ from [¹³C-methyl]naproxen shows that a fraction of naproxen-derived methyl groups is directly metabolized by methylotrophic methanogens, representing a secondary but functionally important route of methyl flux under anaerobic conditions.

### Community shift on caffeine

Demethylation in the caffeine treatments is best explained by archaeal N-demethylation, as the ^13^C-methane is the sole product derived from the N-[^13^C-methyl] group of caffeine, consistent with direct methanogenic demethylation. Although overall caffeine turnover was minimal, several lineages of bacteria and archaea shifted consistently across replicates amended with caffeine, but not in controls. These shifts align with the isotope evidence: with methanogens conducting the primary N-demethylation step, whereas bacteria participate largely in fermentation and syntrophic transformation of downstream caffeine-derived intermediates.

Bacterial taxa enriched under caffeine predominantly belonged to fermentative or syntrophic lineages including: an uncultured class within *Atribacterota* (JS1, enriched 3.5-fold, p≤0.05, Fig. 2A), several *Chloroflexota* groups (*Anaerolineae* g_M2005 sp027036755, 8.7-fold, Fig. 2E.; uncultured *Ardenticatenales* 1.92-fold, p≤0.05, Fig.2A), members of phylum *Pedosphaeraceae* within *Verrucomicrobiota* (enriched 1.9-fold, p≤0.05, Fig. 2A), and a deep sea uncultured Gammaproteobacteria within *Woeseiaceae* (g_QNFK01 sp003645495, enriched 8.7-fold, Fig. 2E). A sulfate reducer within *Desulfobaccia* (g_VGSQ01 sp016874975, enriched 8.7-fold, Fig. 2E), likely persisted on the sulfate in our slurries.

Consistent with the exclusive formation of ^13^CH_4_, archaeal N-demethylators like *Methanosarcina* were enriched (not present in control, Fig. 2E), which are known to encode the N-methyltransferases required to transfer methyl groups to coenzyme M prior to reduction to methane. *Methanosarcina* therefore represent the sole drivers of caffeine-derived N-methyl group turnover.

Caffeine also enriched archaeal lineages that complement methanogenic N-demethylation, by fermenting and syntrophically processing the organic intermediates after degradation. This includes multiple *Bathyarchaeia* (unique to caffeine incubations, Fig. 2E), which can degrade aromatic and complex intermediates, and a member of *Thermoplasmatota* (DHVEG-1 unique to caffeine incubations), which may contribute additional heterotrophic capacity. Fermentation by these archaea, together with bacterial syntrophs, could provide the substrates supporting the primary methanogen in these incubations *Methanosarcina mazei*.

The exclusive production of ¹³CH₄ from ¹³C-caffeine, with no detectable ¹³CO₂, confirms that caffeine-derived N-methyl groups were routed into the methanogenic methyltransferase pathway directly.

Combined with the selective enrichment of methylotrophic *Methanosarcinales*, and the absence of bacterial demethylation signatures (lack of ^13^CO_2_ accumulation and absence of bacterial demethylators) demonstrate that archaea are the primary N-demethylation route of caffeine under methanogenic conditions.

### Bacterial demethylation potential in metagenome-assembled genomes

Because the demethylating potential of novel lineages enriched under micropollutant exposure is unknown, we analyzed bacterial metagenome-assembled genomes (MAGs) for O- and N-demethylation genes.

Methanogenic archaea like *Methanococcoides spp.* and *Methanosarcina spp.*^35,41^ are well established in performing N-demethylation, while rare lineages like *Methermicoccus shengliensis*^42^ and some *Bathyarchaeia* can perform O-demethylation^43^, nevertheless, the recovery of archaeal MAGs was limited, preventing confident reconstruction of key methyltransferase operons (e.g., *mtxA/B/C* and their activators) and precluding reliable pathway-level inference in archaea.

Therefore, we concentrated on bacterial MAGs. We built profile Hidden Markov Models (HMMs) for methyltransferase gene clusters (*MtxA/B/C* and associated activases) using reference sequences from model organisms (*Eubacterium limosum*^28,44^, *Desulfitobacterium hafniense* Y51^45^, *Acetobacterium woodii* ^46^, *Moorella thermoacetica* ^47,48^, *Acetobacterium dehalogenans*^48^). Five clusters were recovered, resolving into groups dedicated to N-demethylation and O-demethylation with only *MtxB* genes, plus mixed N/O-demethylation clusters. The distinct grouping of *MtxB* genes within these clusters is critical, as *MtxB* enzymes are substrate-specific (Fig. 3A). We thus focused on the *MtxB* sequences within the O-demethylation cluster to identify MAGs of anaerobic taxa that possess the genetic potential for naproxen O-demethylation.

**Figure 3.**
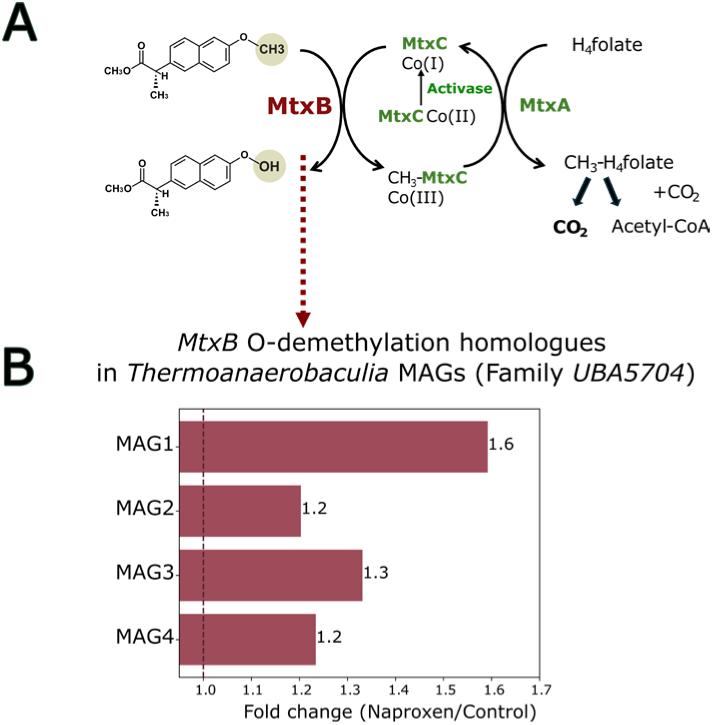
**Enzymatic mechanism for O-demethylation and naproxen enriched anaerobic MAGs with O-demethylation potential**. (A) Proposed biochemical pathway for the anaerobic demethylation of naproxen. The mechanism involves a three-component corrinoid-dependent methyltransferase system: the substrate-specific methyltransferase (*mtxB*) cleaves the O-methyl group from naproxen and transfers it to the corrinoid protein (*mtxC*). The methyl group is subsequently transferred by *mtxA* to tetrahydrofolate (H_4_folate), forming methyl-tetrahydrofolate (CH_3_-H_4_folate), which enters the Wood-Ljungdahl pathway to produce CO_2_ and acetyl-CoA. A reductive activase is required to maintain the corrinoid cofactor in its active Co(I) state. (B) Relative abundance of *mtxB* O-demethylation homologs in four metagenome-assembled genomes (MAGs) of the *Thermoanaerobaculia* family UBA5704. The bar plot displays the fold change of the *mtxB* relative gene abundance in naproxen-treated microcosms relative to unamended controls. Values above the 1.0 threshold (dashed line) show enrichment above controls.

We identified O-demethylation potential in members of the *UBA5704* lineage from the *Acidobacteriota* class *Thermoanaerobaculia* (Fig. 3B). Four distinct MAGs from this lineage exhibited a clear increase in relative abundance in naproxen-treated samples compared to controls, with fold-change values ranging from 1.2 to 1.6. This adds to our observations regarding unknown *Acidobacteriota* ASVs, which were enriched during naproxen degradation (Fig. 2A). Other studies looking at peatland communities identified an O-demethylation potential in *Acidobacteriota* ^49^, suggesting that the capacity for methyl-group turnover extends beyond a single environment and may be an overlooked feature of this group.

### Potential pathway of naproxen degradation

Based on isotopic tracing, taxonomic profiling, and metagenomic analysis, we propose a primary pathway for naproxen degradation in anoxic marine sediments. In contrast, the minimal degradation of caffeine does not support a similarly resolved pathway reconstruction.

The accumulation of desmethylnaproxen, coupled with strong ^13^CO_2_ enrichment (Δδ^13^C = 53 ± 16‰), indicates that bacterial O-demethylation represents the initial and dominant transformation step. This process is likely mediated by taxa such as *Eubacteriales,* known for aryl O-demethylation, as well as *Thermoanaerobaculia (UBA5704)*, whose MAGs encoded abundant O-demethylation genes in naproxen-amended microcosms.

Following O-demethylation, downstream intermediates of naproxen degradation are likely processed via a syntrophic degradation network. This interpretation is supported by the concurrent enrichment of fermentative microorganisms (*Peptostreptococcales-Tissierellales, Paludibacteraceae, Lokiarchaeia),* together with taxa frequently associated with syntrophic metabolism (*Bathyarchaeia, Syntrophomonadaceae*), and the enrichment of hydrogenotrophic methanogens that could serve as H_2_-scavenging partners, such as *Methanogenium cariaci*. The substantial production of ^13^CO_2_ from naproxen, coupled with a nearly identical ^13^CO_2_ production pattern in acetate-amended incubations, further suggests that acetate is a key intermediary metabolite during naproxen degradation (Fig. 4).

**Figure 4.**
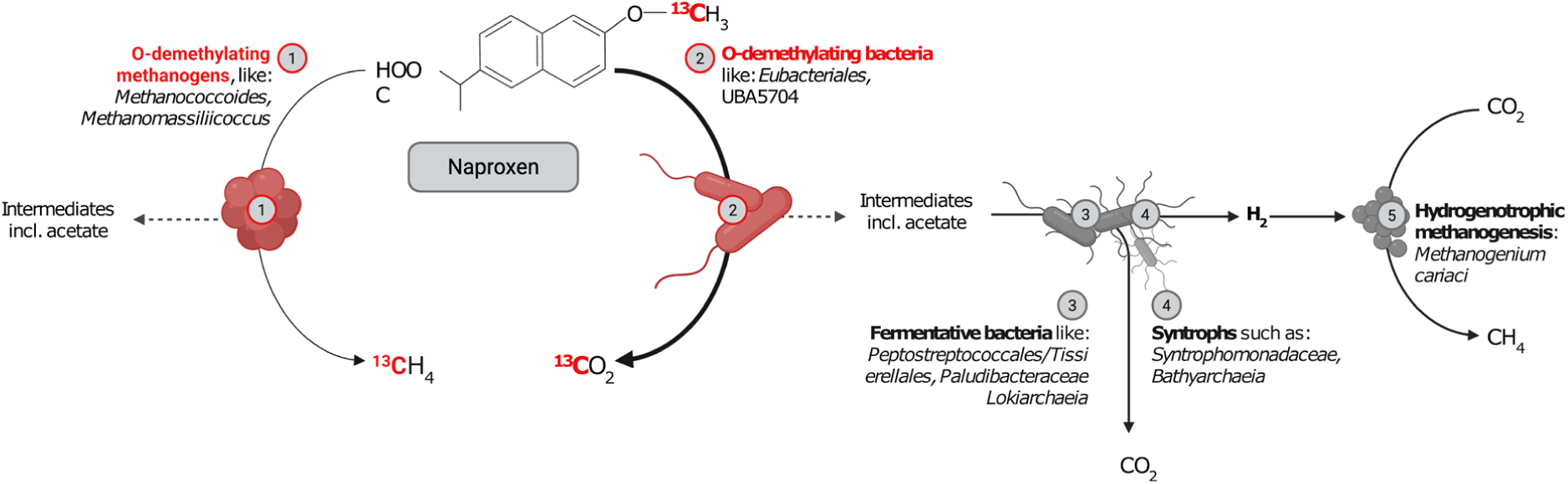
**Conceptual model of anaerobic naproxen demethylation pathways in coastal sediments**. The schematic illustrates the bifurcated routes for the processing of naproxen-derived ^13^C-methyl groups, where relative arrow widths represent the magnitude of carbon flux through the microbial community. In the direct pathway (1), methylotrophic methanogens such as *Methanococcoides* and *Methanomassiliicoccus* perform archaeal demethylation, channeling a fraction of the methyl groups directly into ^13^CH_4_. The primary and quantitatively dominant route (2) is initiated by bacterial O-demethylation, mediated by taxa such as *Eubacteriales* and *Thermoanaerobaculia (UBA5704),* which yields ^13^CO_2_ and organic intermediates. These intermediates, including acetate, are further processed by a network of fermentative bacteria (3), such as *Peptostreptococcales-Tissierellales, Paludibacteraceae,* and *Lokiarchaeia,* and syntrophs (4), including *Syntrophomonadaceae* and *Bathyarchaeia.* This syntrophic degradation generates H_2_ and CO_2_, with the H_2_ subsequently utilized by hydrogenotrophic methanogens (5), such as *Methanogenium cariaci*, to produce CH_4_.

In parallel, a secondary demethylation route appears to involve methylotrophic methanogens (including *Methanococcoides vulcani* and *Methanomassiliicoccales*). The observed enrichment of ^13^C-methane, albeit smaller when compared to ^13^CO₂, indicates that direct methylotrophic methanogenesis from the naproxen-derived methyl group occurs but represents a minor pathway. Collectively, these data support a degradation model initiated by bacterial O-demethylation, in which the methyl carbon is predominantly funneled to CO₂ via fermentative and syntrophic networks, with only a minor fraction diverted to CH₄ via parallel methanogenic pathways (Fig. 4).

Previous studies have established that O-demethylation is the predominant pathway for naproxen degradation in consortia from anaerobic digester sludge and wastewater derived enrichment cultures^18,19^. In sulfate-rich estuarine sediment enrichment cultures, naproxen O-demethylation was associated with a consortium of gut-associated fermenters (e.g., *Prevotellaceae, Campylobacteraceae*) and acetogens (e.g., *Lachnospiraceae, Eubacteriaceae*), though the subsequent fate of the removed methyl group has not been thoroughly investigated^19^. Our findings extend this work by showing that, naproxen-derived methyl groups are ultimately mineralized to CO_2_ and CH_4_, through the combined action of, fermenters, syntrophs and hydrogenotrophic and methylotrophic methanogens.

Methyl groups represent one of the most common substituents in pharmaceuticals and pesticides^50^ ensuring that methylated compounds form a substantial portion of the complex contaminant mixture discharged from WWTPs into aquatic systems^2^. The persistence of caffeine, contrasted with the active transformation of naproxen, highlights an important environmental implication for sediments receiving wastewater effluents. Chronically micropollutant-exposed sediments appear ineffective at removing the micropollutants, particularly after burial in anoxic zones where these can accumulate over long timescales^51^. Although methane production from any individual micropollutant is small, the continuous local influx of diverse methylated compounds from pesticides and pharmaceuticals may represent an overlooked carbon source for coastal methanogenesis. Future studies should therefore move beyond single-compound assessments and evaluate how structurally diverse methylated micropollutants, occurring as complex mixtures, influence sedimentary methane cycling and greenhouse gas dynamics.

Overall, this study reveals that the molecular structure of methylated pollutants is a key factor determining their microbial fate in coastal sediments, which in turn governs their impact on methane production. Through integrated isotopic tracing, metabolite analysis, and functional metagenomics, we resolved the pathway of methyl group processing: naproxen’s O-methyl group enters syntrophic bacterial networks (e.g., *Eubacteriales*, *Thermoanaerobaculia*) yielding predominantly CO₂, with only a trace diverted to methane. In contrast, caffeine’s recalcitrant N-methyl groups feed directly into archaeal methanogenesis (*Methanosarcina*, *Bathyarchaeia*), also at low levels due to the compound’s slow breakdown. Consequently, these findings provide the mechanistic, process-oriented framework required to predict how the diverse, wastewater-derived methylated compounds entering coastal environments will differentially sustain microbial activity and influence greenhouse gas fluxes.

## Methods

### Sample collection, biogeochemical, and micropollutant analysis

Sediment cores were collected in August 2022 in the central part of Hakefjorden fjord, approximately 1 km off the town of Stenungsund (Swedish West Coast). This sampling site was selected based on previous studies reporting high concentration of micropollutants in the Hakefjorden fjord^6^. The sampling site was 50 m north of a marine outflow discharging treated wastewater from the Strävliden WWTP (coordinates: 58.09553° N 11.8061° E). The sampling location had a water depth of 13.3 m, with the bottom water exhibiting a salinity of 28.4 ppt and a temperature of 16.3°C. Three sediment cores were collected and sliced onboard R/V Alice. The upper layer, ranging from 2 to 10 cm in depth, was stored in glass bottles and kept at 4 °C, under an anoxic (N_2_) atmosphere for 4 days until microcosm experiments.

Sediment geochemical parameters were determined through depth including methane (CH_4_), dissolved inorganic carbon (DIC), sulfate, and other nutrients (nitrate, ammonia, phosphate). Methane samples were collected at 3 cm intervals using 3 mL cutoff syringes via predrilled and taped holes, then transferred to 21 mL vials containing 12 mL of 5 M NaCl and 0.0241 M ZnCl₂ to inhibit microbial activity, after which the vials were sealed. The CH_4_ concentrations were determined using gas chromatography^52^, with the method quantification limit (LOQ) for methane at 1.8 ppm. Furthermore, DIC, sulfate and nutrients from porewater samples were collected using rhizons (Rhizosphere Research Products, 0.45 µm pore size), which were inserted at 2 cm intervals in predrilled holes. The collected porewater was either stored in exetainers and refrigerated for DIC analysis or placed in 1.5 mL centrifuge tubes and frozen with the addition of 20 % sodium acetate for sulfate analysis. Sulfate concentrations were measured via ion chromatography, as previously described^53^, with the LOQ for all ion chromatography measurements at 10 µM. Nutrient analyses were performed spectrophotometrically according to established protocols^54,55^.

DIC concentration analysis was performed using a custom-made gas stripping system with N_2_ as the carrier gas and a Li-COR 6262 analyzer. Samples were acidified with phosphoric acid (8.5 %) to convert carbonate species into CO_2_ gas, which was measured using non-dispersive IR spectrometry^56^. Sediment porosity was determined by calculating the ratio between the wet weight and dry weight of the sediments, where the dry weight was measured by drying the sediments in an oven at 30°C until a constant weight was achieved. Organic carbon content, *f_OC_*, was determined with an elemental analyzer isotope ratio mass spectrometer (IRMS), after removing inorganic carbon via acidification with hydrochloric acid^57^.

Micropollutant profiles were built based on a target list of 861 organic micropollutants. Micropollutants were extracted from sediment samples by pressurized liquid extraction (PLE) with a methanol:0.1 M citrate buffer:0.1 M NaEDTA solvent mixture (60:30:10) followed by solid-phase extraction (SPE) for enrichment and cleanup, and then analyzed using liquid chromatography-high resolution mass spectrometry (LC-HRMS), with LOQ for LC-MS pollutants at 5 nM^6^.

### Slurry incubations

Slurries were prepared by mixing homogenized sediments extracted from three cores (depth range: 2-10 cm) with bottom seawater (collected using a Niskin bottle). The 20 mL slurry was added to 120 mL vials pre-filled with 30 mL of a modified DSM 141c marine methanogenic media lacking sodium acetate, yeast, trypticase peptone, trimethylammonium chloride, and resazurin. This entire process was conducted under an anaerobic atmosphere (N_2_:H_2_, 98%:2 %) in a COY glovebox where the glass vials were closed with blue butyl-rubber stoppers. Afterwards the headspace was flushed with a CO_2_:N_2_ gas mixture (20:80, v/v) to remove traces of H_2_. Each serum vial was amended with one of the five ^13^C-labeled compounds, namely: 10 mM sodium acetate-2-^13^C (^13^CH_3_COONa), 10 mM sodium ^13^C-bicarbonate (NaH^13^CO_3_), 10 mM ^13^C-methylamine-hydrochloride (^13^CH_3_NH_2_ · HCl), 0.5 mM (±)-naproxen-(methoxy-^13^C) (^13^CC_13_H_14_O_3_) and 0.5 mM caffeine-(trimethyl-^13^C_3_) (^13^C_3_C_5_H_10_N_4_O_2_). Microcosms were started over a two-day period, with two sets of triplicate unamended controls incubated each day. For the carbonate treatment, the microcosm headspace was flushed with H_2_:CO_2_ (20:80), and subsequent flushing occurred on day 12 to counter the negative pressure inside these bottles. This is expected to remove some of the labeled ^13^CO_2_ that has equilibrated with the headspace atmosphere. Unless otherwise stated, all treatments were performed in four replicates at a constant temperature of 25 °C in the absence of light and agitation, for 24-25 days.

### Sampling and analytical procedures

Samples to determine carbon isotope signature (δ^13^C_CH4_, δ^13^C_DIC_), sulfate, acetate, naproxen, desmethylnaproxen and caffeine concentrations were collected using hypodermic needles and syringes preflushed with N_2_:CO_2_ at the beginning and the end of the incubation period. Methane samples were taken over the course of the incubations at the start (day 0) and on days 1, 4, 12, and 24 for caffeine, acetate, carbonate, methylamine, and control treatments. For naproxen and the second series of control incubations, methane concentrations were analyzed at the beginning (day 0) and on days 1, 6, 13, and 25. CH_4_ concentrations were measured by gas chromatography, and sulfate and acetate concentrations were measured by ion chromatography as previously described^53^.

Samples for δ^13^C_DIC_ were carefully filled to the brim to prevent gas bubbles from forming in 2-mL HPLC glass vials, each containing 20 µL of HgCl_2_-saturated water. The conversion of DIC to CO_2_ was achieved through acidification using 500 µL of 40% H_3_PO_4_ and then equilibrated in the headspace overnight at 70 °C inside a He-flushed 10-mL vial. Subsequently, δ^13^C_DIC_ isotope ratios were measured with IRMS, as previously described^58^. For δ^13^C_CH4_ analysis, a volume of 1 mL of headspace gas from the microcosm bottles was carefully transferred into 12-mL exetainers, which were previously flushed with a CO_2_:N_2_ gas mixture (20:80, v/v) and contained 6 mL of 0.5M NaCl and 1 mL of 20 % ZnCl_2_. After subsampling, the exetainers were stored upside down before analysis at the Stable Isotope Facility at the University of California, Davis. The samples were analyzed using a preconcentration unit interfaced with an IRMS, as previously described^59^. Samples from carbonate treatments were not analyzed for isotopes of methane and DIC as this is the primary methanogenesis pathway in the marine environment.

Organic micropollutants (caffeine, naproxen, and desmethylnaproxen) from the microcosm experiment were extracted using liquid-liquid extraction. Since naproxen degraded substantially, we measured both naproxen and desmethylnaproxen, while caffeine was measured directly due to minimal degradation. In brief, 0.5 mL of media from the microcosm was extracted three times with 1 mL of ethyl acetate, employing vortexing, centrifugation, and subsequent collection of the organic phase each time. The combined ethyl acetate phase was then evaporated to dryness, and the resulting residue was reconstituted in 0.5 mL of methanol before being analyzed using LC-HRMS, following the previously described procedure^6^. The total quantity of the chemical present per bottle was determined by utilizing the carbon-water partition coefficient obtained from EPI Suite™ and the measurement of organic carbon content in the sediment.

### Microbial community analysis by amplicon sequencing and metagenomics

*16S rRNA.* Sediment samples were collected in triplicates for microbial metabarcoding community characterization from each of the treatments at the conclusion of the incubation period. Additionally, to best represent sediment communities in their initial state, we collected duplicate samples of fresh slurries from unincubated control (original sediment), which was not subjected to incubation.

All sediment samples were promptly frozen and shipped to Novogene Europe for subsequent DNA extraction and amplicon sequencing. Modified primers designed to target and amplify the V4-region of the 16S rRNA gene sequence were employed for this purpose^60,61^. Seventeen libraries were sequenced on a paired-end Illumina platform, generating 250bp paired-end raw reads. Sequencing libraries were subjected to rigorous quality control and pre-processing. FastQC software v0.11.9^62^ was used to quality control checks on raw sequences. Cutadapt v3.3^63^ was utilized to eliminate short sequences ”--minimum-length 20” and fastp v0.23.1^64^ to remove low-quality reads using as follows: “-q 19 -u 15”. Chimera removal was performed using VSEARCH 2.16.0^65^. Denoising and clustering into amplicon sequence variants (ASVs) were performed using the denoise-paired method of the DADA2 package v1.14^66^, executed through the bioinformatics platform QIIME2 v2023.5.1^67^. The parameters used to run the denoise-paired method were as follows: p-trunc-len-f 210, p-trunc-len-r 205, p-trim-left-f 22, p-trim-left-r 22, p-max-ee-f 3, p-max-ee-r 3, and p-chimera-method consensus. After this process, more than 80% of the reads were retained, on average. The taxonomy assignment of ASVs was accomplished using the feature classifier method, classify-sklearn, in QIIME2. The SILVA database 138.1 was used as reference^68^, utilizing the pre-trained classifier designed for the 515F/806R region in the Silva database 138 (Silva 138 SSURef NR99 515F/806R region sequences).

To refine the final ASV table, singletons were removed, and the ASV data were classified into bacteria and archaea. To assess the statistical differential abundance of microbial community matrix abundances were used as input for MaAsLin2 v1.8.0^69^ and EdgeR v4.7.0^70^, respectively, within the R environment (ver. 3.6.3)^71^. This function fitted a linear model to the transformed abundance of each feature on the specified sample grouping, assessed significance using a Wald test, and provided Benjamini-Hochberg (BH) FDR-corrected p-values. We applied arcsine square root transformation and total sum scaling normalization, considering only results with FDR-corrected p-values < 0.05.

*Metagenomic analysis.* At the end of the incubation period, triplicate sediment samples from each treatment were homogenized into a single composite sample for metagenomic analysis, ensuring a representative functional profiling of microbial communities. For unincubated controls (original sediment), duplicate fresh slurry samples were homogenized and analyzed to capture the baseline microbial state prior to incubation. DNA was extracted using the DNeasy PowerSoil Pro Kit (Qiagen, Germany), and its quality was assessed using both a Qubit 4 Fluorometer and a NanoDrop OneC Spectrophotometer. Fragment size distribution was evaluated on the Agilent 4200 TapeStation.

Sequencing libraries were prepared with the SQK-NBD114.96 kit (Oxford Nanopore Technologies) and sequenced on a PromethION P24 system using FLO-PRO114M (R10.4.1) flow cells and MinKNOW software version 24.02.19. Basecalling was performed with Dorado v7.3.11 using the super-accuracy model. Reads were quality-filtered using nanoq v0.10.0, retaining sequences with a minimum Phred score of 15 and a minimum length of 1000 bp, while trimming 150 bp from each end.

Assembly was carried out with Flye v2.9 in metagenome mode, and consensus sequences were polished using Medaka v1.11.3. Contigs shorter than 1000 bp were discarded. Reads were aligned back to assemblies using Minimap2 v2.28, and processed with SAMtools v1.20. MAGs were recovered using SemiBin2 v2.1.0, dereplicated with dRep v3.5.0 under thresholds of ≥50% completeness and ≤10% contamination and assessed for quality using CheckM2 and GUNC v1.0.5. Coverage and abundance were estimated with CoverM v0.6.1, applying identity and alignment thresholds of 95% and 90%, respectively. Taxonomic classification of MAGs was performed using GTDB-Tk v2.4.0 with the GTDB R220 reference. Additional taxonomic profiling was conducted with Kaiju v1.9.2 using the nr_euk database, and marker-based classification was carried out with MELON using default parameters. Ribosomal RNA genes were extracted and mapped to the SILVA 138.1 database for classification. Genome annotations were performed with Bakta v1.9.4 for bacterial MAGs and Prokka v1.14.6 for archaeal MAGs.

*Identification and analysis of candidate of N- and O-demethylation genes.* To identify putative genes involved in N- and O-demethylation, a multi-step bioinformatic pipeline was implemented. Previously characterized genes implicated in demethylation were selected as a reference based on previous report^49^.

The protein sequences of the reference genes were used as a query in a BLASTp search against the NCBI non-redundant (nr) protein database. The search aimed to identify homologous proteins across diverse taxa, and top hits were selected based on a minimum sequence identity of 30%, full-length alignment coverage, and the lowest E-values. The retrieved protein sequences were aligned using MAFFT to ensure accurate multiple sequence alignments^72^. These alignments were then used to construct hidden Markov models (HMMs) with HMMER^73^. The resulting HMM profiles were employed to search for candidate demethylation-related proteins within a set of query proteins. HMM search results were filtered to retain only hits with an E-value threshold of ≤1e-10. To explore the relationships between the identified proteins, a network analysis was conducted using R environment. Protein connections were inferred based on pairwise similarity, and networks were visualized using standard R plotting functions.

### Statistical Analysis

Microbial community differential abundance was analyzed using MaAsLin2 and EdgeR, with significance determined at an FDR-corrected p-value < 0.05. Geochemical comparisons between treatment groups were performed using two-sample t-tests (p < 0.05). Data are presented as mean ± standard deviation (SD). Exact biological replicates (n) for each experiment were as follows: all microbial incubations, n = 4 per treatment; geochemical measurements, n=4 per treatment; isotope analyses, n=3 per treatment; 16S rRNA sequencing, n = 3 per treatment; metagenomic composites, n = 1 per treatment.

## Acknowledgements

We thank the R/V Alice staff and Lok Shan Cheung for their assistance during sampling at sea. This work was supported by the Marie Skłodowska-Curie Actions (Project ID: 101062895) and the Swedish Research Council VR (Project ID: 2022-04710). AER and KA are supported by an ERC Consolidator grant awarded to AER, grant number: 101045149.

## Author contributions

T.G., A.-E.R. and S.B. conceived and designed the study. T.G., S.B. and A.F. performed the experiments. T.G., S.K. and M.K. conducted chemical analyses. A.F., K.A. and P.A.I. analyzed sequencing data. T.G. wrote the manuscript with input from all authors. All authors reviewed and approved the final manuscript.

## Data Availability

The raw sequence data from both the 16S rRNA and metagenomic libraries have been deposited in the NCBI Sequence Read Archive (SRA) under the accession number PRJNA1335822. The micropollutant screening data are available on Zenodo (10.5281/zenodo.15268726). All other data supporting the findings of this study are available within the article and its Supplementary Information, or from the corresponding author upon reasonable request.

